# Defining the DmsA Signal Sequence Interaction with DmsD And TatBC

**DOI:** 10.1101/789909

**Authors:** Deepanjan Ghosh, Sureshkumar Ramasamy

**Affiliations:** From the Division of Biochemical Sciences, CSIR-National Chemical Laboratory, Pune-411008, India

## Abstract

Redox – enzyme maturation proteins (REMP) ensure co-factor loading and folding of proteins targeted to the twin-arginine translocation (Tat) pathway. The details of the interaction of a REMP with the corresponding signal sequence of its substrate are not well understood. Here, we demonstrate the features of the signal sequence for the Tat substrate DmsA (ssDmsA) responsible for complex formation with its REMP, DmsD, and with the Tat membrane complex TatB & TatC (TatBC). A heterologously expressed ssDmsA/DmsD complex forms two stochiometric populations corresponding to monomeric and dimeric forms of the complex. The monomeric complex has a higher affinity for the TatBC complex than the dimeric, which imply higher level regulation process to ensure the maturation of protein before translocation. Results from various binding studies yielded the shortest signal peptide required for ssDmsA/DmsA interaction and the region responsible for the TatBC interaction. Further experiments like alanine scanning in this peptide highlight the possible residues that are essential for this complex formation.

---

The twin-arginine translocation (Tat) pathway is responsible for the post-translational transport of folded proteins across the cytoplasmic membrane of bacteria, archaea and the thylakoid membrane of the plant chloroplast. The minimal Tat system, here described for *E. coli*, is composed of the integral membrane proteins TatA, TatB and TatC. Substrates of this pathway contain a characteristic Tat specific Arg-Arg motif in the N-terminal signal sequence (1). The Tat signal localizes the protein to the membrane where it is recognized by TatC (2) and then the protein is translocated across the cytoplasmic membrane though a pore formed by TatA (reviewed in (3–5)). In most cases, Tat substrates are either a component of the respiratory electron transport chain or are bacterial virulence factors (6,7). These substrates often incorporate large co-factors or assemble into complexes before getting translocated (8).

Dimethylsulphoxide reductase is a heterotrimeric periplasmic complex that uses dimethylsulphoxide (DMSO) as an electron acceptor during anaerobic respiration. The central protein, DmsA, is synthesized in the cytoplasm, loaded with its catalytic molybdopterin (MoPt) co-factor and the accessory proteins DmsB and DmsC. This enzyme complex is an example of Tat substrates that must be fully assembled prior to translocation to the bacterial periplasm (9). Delivery of these proteins is monitored by a special class of cytoplasmic chaperones that bind specifically to the Tat signal of their substrate protein masking the signal sequence thereby ensuring proper maturation and co-factor loading (10,11). These chaperones, dubbed redox-enzyme maturation proteins (REMP), prevent the futile export of immature protein (12). Each REMP generally has a specific binding partner and there are a number of verified examples. DmsD, a well characterized REMP, binds the DmsA N-terminal Tat signal sequence which contains the Tat consensus motif (S/TRRxLVK) (13) (14).

Generally signal peptides have tripartite structure with a positively charged N-region, a middle hydrophobic stretch followed by a polar C-region that contains a signal peptidase cleavage site (15,16). The *E. coli* genome encodes at least 29 putative signal sequences that contain the Tat motif. (17). Many studies have emphasized the significance of the presence of RR in the signal sequence during translocation (2,18,19). The Tat signal is targeted to the pre-formed complex between TatB and TatC, the TatBC recognition complex (20,21). Replacing the RR of the signal sequence with a pair of lysines retains the ability to co-purify with the TatBC complex. This suggests that the essential RR residues are not necessary for binding to the TatBC recognition complex but are required for sucessful entry into the transport cycle (22). Additional evidence revealed a major role for TatB in initial binding of the precursor protein (22). The interaction between TatBC and the Tat signal of DmsA has been captured by bimolecular fluorescence complementation (BiFC) (23). The residues essential for the TatC recognition has been studies extensively (19) Random mutational analysis in the Tat signal peptide indicated that the some of gain of function mutation with respect to rate of translocation occurred within or near the Tat motif, highlighting the significance of this region (24).

Other characterized REMP/signal peptides are TorD/TorA (8) NapD/NapA (25), NarJ/NarG, NarW/NarZ and HybE/HybO (26). Other chaperones, such as DnaK and SlyD, were shown to bind broad range of different Tat signal sequences (27). Bindings of these chaperones are not essential for the Tat depended transport of substrate across the membrane (28,29).

The role of the signal sequence is more than a simple targeting factor. This is exemplified by the fact that substitution of a DmsA signal with a TorA signal sequence peptide showed poor growth in anaerobic condition despite the targeting of DmsABC to the membrane (9). Without a correct signal sequence the maturation and production of functional enzyme prior to targeting is lost indirectly implying the significance of DmsD binding for the production of functional enzyme (9). DmsD can form a complex with the N-terminus of DmsA independent of the globular domain of the protein (13). *In vivo* studies suggest that DmsD is not required for targeting (29). Also there is evidence that signal peptide is extensively crosslinked to ribosomal components and the trigger-factor chaperone during synthesis but does not play a critical role in the export of Tat dependent protein (30)

Despite structural information, the molecular characteristics of REMP/signal sequence binding have not been fully elucidated (31) (32). REMP chaperones contain two specific motifs Y/F/W-X-X-L-F and E-(P-X/X-P)-D-H/Y that are conserved among all members of this family and mutations in these regions lower the binding affinity to substrates (33).

The binding of REMPs to signal sequences has been localized to the Tat signal and the possible protease site (23). Further studies using DmsD and the signal peptide indicate binding via a hydrophobic interaction with micromolar affinity in an equimolar ratio (34). More extensive work has been done on the interaction between TorD and the signal peptide of TorA. Here, recognition has been localized to the hydrophobic h-region (35). Glutamine-scanning mutagenesis through this region demonstrated that an L31Q variant of the TorA signal peptide impaired binding of TorD (36). Synthetic peptide truncation variants of the TorA signal showed weaker or no binding of TorD with C-terminal truncations while N-terminal truncations, including the twin arginine motif, did not dramatically affect TorD binding (35). This suggests that the twin-arginine motif is not essential for chaperone recognition.

Many mechanistic questions about the role of REMPs remain. The coordination between co-factor loading and the subsequent transfer of the mature protein to the translocase assembly is unknown.

Our specific goals in this study were the biochemical characterization of DmsD and the signal sequence of DmsA (ssDmsA) complex, also the complex with translocase assembly and to map the location of the binding site of TatBC and DmsD in the DmsA signal sequence, thereby deriving the smallest length of sequence essential for a specific interaction.

## EXPERIMENTAL PROCEDURES

### Cloning, Over-expression and protein purification

The gene encoding *Ec* ssDmsA was amplified from *E. coli* genomic DNA using the primers 5’-GGGGGC**CCATGG**GCAAAACGAAAATCCCTGATGCGG-3’ and 5’-GGGGCG**GCTAGC**AATGGCGCTATCGACAGCG-3’ which incorporate the restriction endonuclease sites NcoI and NheI (highlighted in bold), respectively. Digested PCR product was ligated into a modified T7 promotor-based pET33b expression vector (Novagen) that contained the super-folding variant of green fluorescent protein encoding gene with C-terminal 6xHis-tag and a modified multiple cloning site. The truncation mutants of ssDmsA obtained by designing appropriate primers for quick-change method. Site-specific Ala mutants for all the residues and Ser mutation for Ala residues were introduced into ssDmsA using the quick-change method (37). All clones obtained from PCR-amplified DNA were sequenced to ensure that no undesired mutation has been introduced.

The plasmids were transformed into *E*.*coli* BL21 (DE3) Tat mutant cells. The plated cells were directly inoculated into 1 L of LB medium containing 35 µg ml^−1^ kanamycin. The culture was grown at 310 K to an OD_600_ of 0.8, followed by induction with 300 µM isopropyl β-D-1-thiogalactopyranoside (IPTG from Anatrace) for three hours. The cells were harvested by centrifugation and the cell pellet was resuspended in 100 ml of Buffer A (100 mM NaCl, 10 mM β-ME, 30 mM imidazole and 50 mM Tris HCl pH 7.5) then lysed by passing three times though a microfluidizer M-110L (Microfluidics). Cell debris was removed by centrifugation at 18,000g for 30 minutes at 277K. The supernatant was loaded onto gravity column containing 5 ml Ni-NTA affinity resin (Qiagen) pre-equilibrated and washed with 30 ml of Buffer A then eluted with Buffer A containing 300mM imidazole. The purified protein was concentrated to 15 mg ml^−1^ using a centrifugal concentration device (Amicon ultra, Millipore) before storage at 193K. The DmsD and DmsD-MBP fusion protein were cloned and purified as described previously (31). The *Ec*TatB and *Ec*TatC were cloned in the pACYC duet system (pACYCDuet-1) (Invitrogen) with and without tag. The purification procedure has been followed as given in elsewhere (38).

### Affinity Capture

The affinity capture of a partner either *in vivo* or *in vitro* performed by co-expressing DmsD without tag and ssDmsA with His-tag (*in vivo*) in BL21 (DE3) Tat mutant cells. For *in vitro* analysis saturated amount purified DmsD-MBP fusion protein was mixed with ssDmsA-GFP-6xHis protein and incubated at 277K for 1 hr. For *in vivo* analysis, co-expressed cells were harvested after 3 hrs of induction, mixed with 1 ml of Buffer A and sonicated. The supernatant was passed though the Ni resin in a spin colum which was then washed thoroughly with Buffer A containing 30mM imidazole and eluted in 100 µl of BufferA containing 300mM imidazole. The association of the DmsD is confirmed by SDS-PAGE (*in vivo*) or by passing through the SEC (*in vitro*).

To check the interaction between the DmsD/ssDmsA complex and the *Ec*TatBC assembly, the over expressed cells of *Ec*TatBC without 6xHis tag were lysed by passing four times though a microfludizer and the total membrane fraction was isolated by centrifugation at 45,000g at 277K for 1 hr. The membrane pellet was solublized in 1% digitonin, centrifuged at 35,000g at 277K for 30 min. The supernatant was used for the pull-down experiments; the supernatant was diluted 10 fold with the buffer A, to minimize the cause of excess detergent in complex formation.

### SPR analysis

All SPR analysis was performed at 298K on a Biacore T100 system using Biacore CM5 research grade sensor chips (GE Healthcare). Purified fractions of DmsD were immobilized on sensor chips using amine coupling at 400 μg/ml of total protein. The dextran matrix on the sensor chip surface is first activated with 1:1 mixture of 0.4 M 1-ethyl-3-(3-dimethylaminopropyl) carbodiimide (EDC) and 0.1 M N-hydroxysuccinimide (NHS) to create reactive succinimide esters. The maximum immobilization level of the DmsD was achieved after three injections and the sensogram baseline was increased to 2000 RU. The surfaces were deactivated by passage of ethanolamine-HCl (1.0 M at pH 8.5). Experiments were carried out in running buffer containing 10 mM HEPES pH 7.4, 150 mM NaCl, and 0.005% v/v Surfactant P20 (GE Healthcare). Each of the ssDmsA mutants was analyzed for binding by injecting 200 μl of a serially diluted (10, 20, 40, 80, 160 and 320 nM) purified sample at a flow rate of 10 μl/min for 10 minutes. The dissociation step was performed at the same flow rate with same buffer for 15 minutes. Regeneration solution (1 M NaCl, 100 mM glycine pH 9.5) for 2 minute at 5 μl/min were used to regenerate the DmsD surface after each injection. For the *Ec*TatBC complex, purified fractions of *Ec*TatBC with 6xHis tag was used directly for immobilization on sensor chips using the amine coupling in presence of 0.1% (w/v) digitonin at 250 μg/ml of total protein. The dextran matrix on the sensor chip surface was prepared as mentioned above. The maximum immobilization level of the *Ec*TatBC was achieved after three injections and the sensogram baseline was increased to 2000 RU. The surfaces were then deactivated by passage of ethanolamine-HCl (1.0 M at pH 8.5). Experiments were carried out in running buffer containing Sodium acetate buffer pH 5.0 with NaCl 100mM, 10% v/v glycerol and 0.1% (w/v) digitonin. Each of the ssDmsA mutants, DmsD and ssDmsA/DmsD complexes were analyzed for binding by injecting 200 μl of a serially diluted (100, 200, 400, 800, 1600 and 3200 nM) purified sample in PBS buffer contain 10% v/v glycerol and 0.1% (w/v) digitonin at a flow rate of 10 μl/min for 10 minutes. The dissociation step was performed at the same flow rate with same buffer for 15 minutes. Regeneration solution [1 M NaCl, 100 mM glycine pH 9.5 contains 10% (v/v) glycerol and 0.1% (w/v) digitonin] for 2 minutes at 5 μl/min were used to regenerate the *Ec*TatBC surface after each injection.

One flow cell of the CM5 chip was used as the blank control. The signal of each binding experiment was corrected for non-specific binding by subtracting the signal obtained for a blank surface. The kinetic of the association and dissociation phase were measured at several flow rates from 5 to 40 μL. the kinetic rates measured were not affected by flow rate, demonstrating that the system is not mass-transfer limited. Kinetic data analysis was done in BIAevaluation software v. 4.1 for multiple concentrations simultaneously using a 1:1 Langmiur model. Some data curves were eliminated to reduce the χ2 value for a better fitting; A more detailed description of the parameters and equation is provided in the Biacore (GE Healthcare) T100 software Handbook (BR-1006-48, edn AE, 186-187)

### CD analysis

All CD measurements were performed on a JASCO J-715 spectropolarimeter, using circular quartz cells with path lengths of 0.02 cm. The response time setting for the spectrometer was 2 s with a data acquisition time of 5 s. The CD spectra are reported in terms of [e], molar ellipticity, deg cm^2^ dmol. Each measurement was the average of three repeated scans in steps. The temperature of the sample was controlled by a circulating water bath (Lauda, type K2R) linked to the outer jacket of the cuvette (Helmann).

For CD measurements of the complex of DmsD and ssDmsA peptides, a molar ratio of 1.5:1 (peptide:protein), at a peptide concentration of 50 µm, was used, which is comparable to the K_D_, value of DmsD to the peptides. The differential spectrum of DmsD was obtained by subtracting the CD spectrum of the peptides from the CD spectrum of the complex at the same concentration as in the complex before the conversion to the ellipticity values.

### Multi-angle light-scattering analysis

Purified *Ec* ssDmsA and DmsD complex (15 mg ml^−1^) was loaded onto a Shodex protein KW-803 size-exclusion column equilibrated with 50 mM Tris-HCl pH 7.0, 150 mM NaCl, 10mM β-mercaptoethanol and connected in-line with a Dawn 18-angle light-scattering detector coupled to an Optilab interferometric refractometer and a WyattQELS Quasi-Elastic Light-Scattering instrument (Wyatt Technologies). Data analysis was performed with the ASTRA V5.3.4.14 software (Wyatt Technologies) and molecular weights were calculated using the Zimm fit method.

## RESULTS

### Initial characterization of DmsD/ssDmsA complex

The signal sequence of DmsA (first 50 amino acids) was cloned into an expression construct followed by a protease site (Tev), green fluorescent protein (GFP) and a six histidine-tag for purification (ssDmsA-GFP). An expression construct of full length maltose binding protein (MBP) tag followed by a second protease site (Thr) and DmsD (MBP-DmsD). Both native proteins and their mutants were purified to homogeneity by chromotagraphy. Wild type and mutant variants of ssDmsA were purified and tested for their binding capability with wild type DmsD by incubating purified MBP-DmsD with ssDmsA-GFP fusion protein and separating on a size-exclusion column (SEC). The two different purification tags were added for ease of purification of homogenous complex (Figure 1A). The observed results are presented in the figure 2. The SEC profiles clearly show DmsD/ssDmsA complex formation. The formation of complex was further analyzed by mixing different molar ratios of each component. A two-fold excess of ssDmsA-GFP saturates DmsD and forms homogeneous complex DmsD (Figure S1).

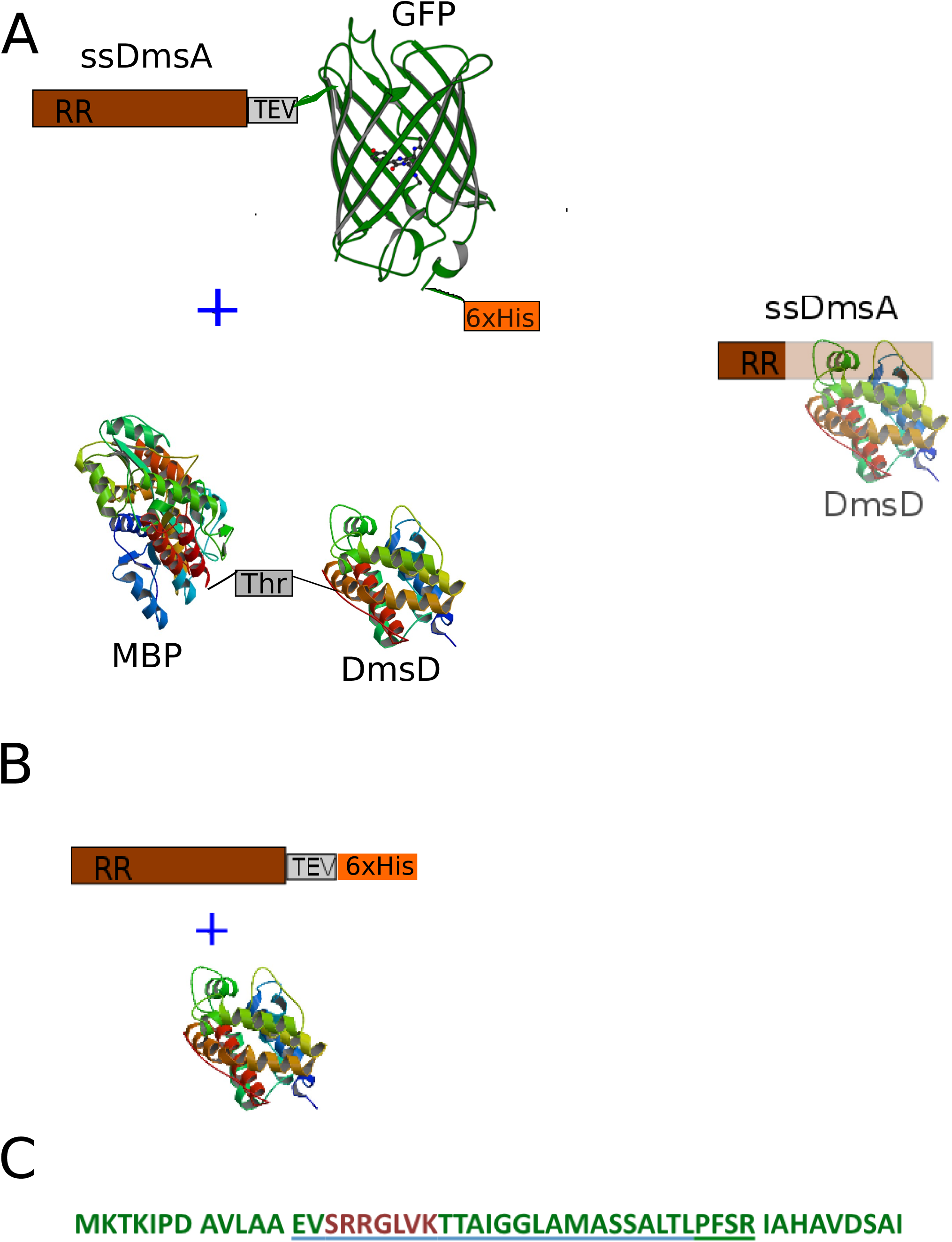

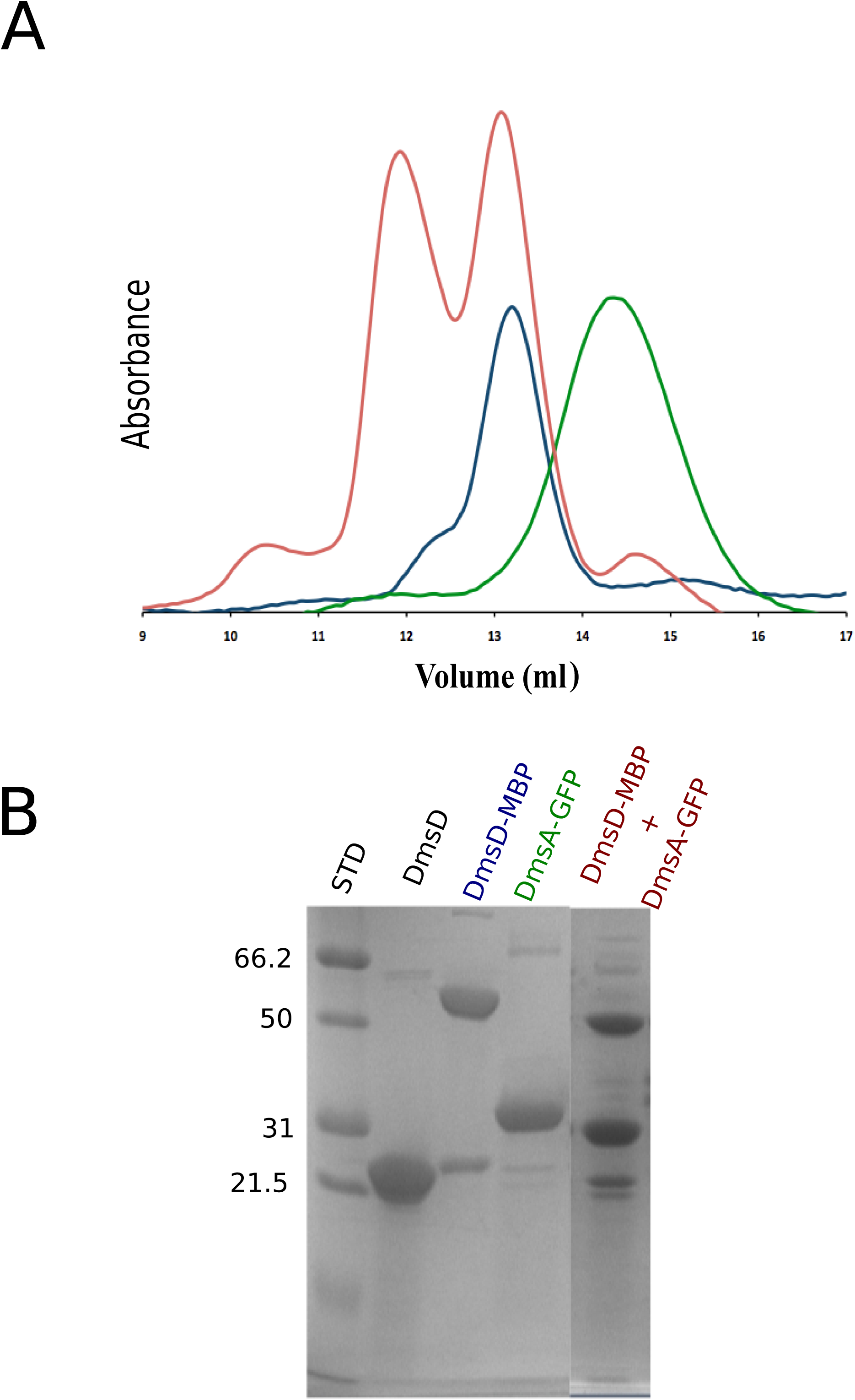

The complex formed between the co-expressed ssDmsA (6xHis tag) with DmsD (Figure 1B) had two distinct populations (Figure 3A), confirmed by SDS PAGE (Figure 3B). A comparison of the elution profiles of monomeric ssDmsA/DmsD complex and DmsD alone are shown in Figure S2. To determine the stoichiometry of the complex we performed multi-angle light scattering (MALS) (Figure S3). The results suggest that DmsD/ssDmsA complex exists either as a single copy of ssDmsA and DmsD (monomeric) or one copy of ssDmsA and two copies of DmsD (dimeric).

**Figure 3.**
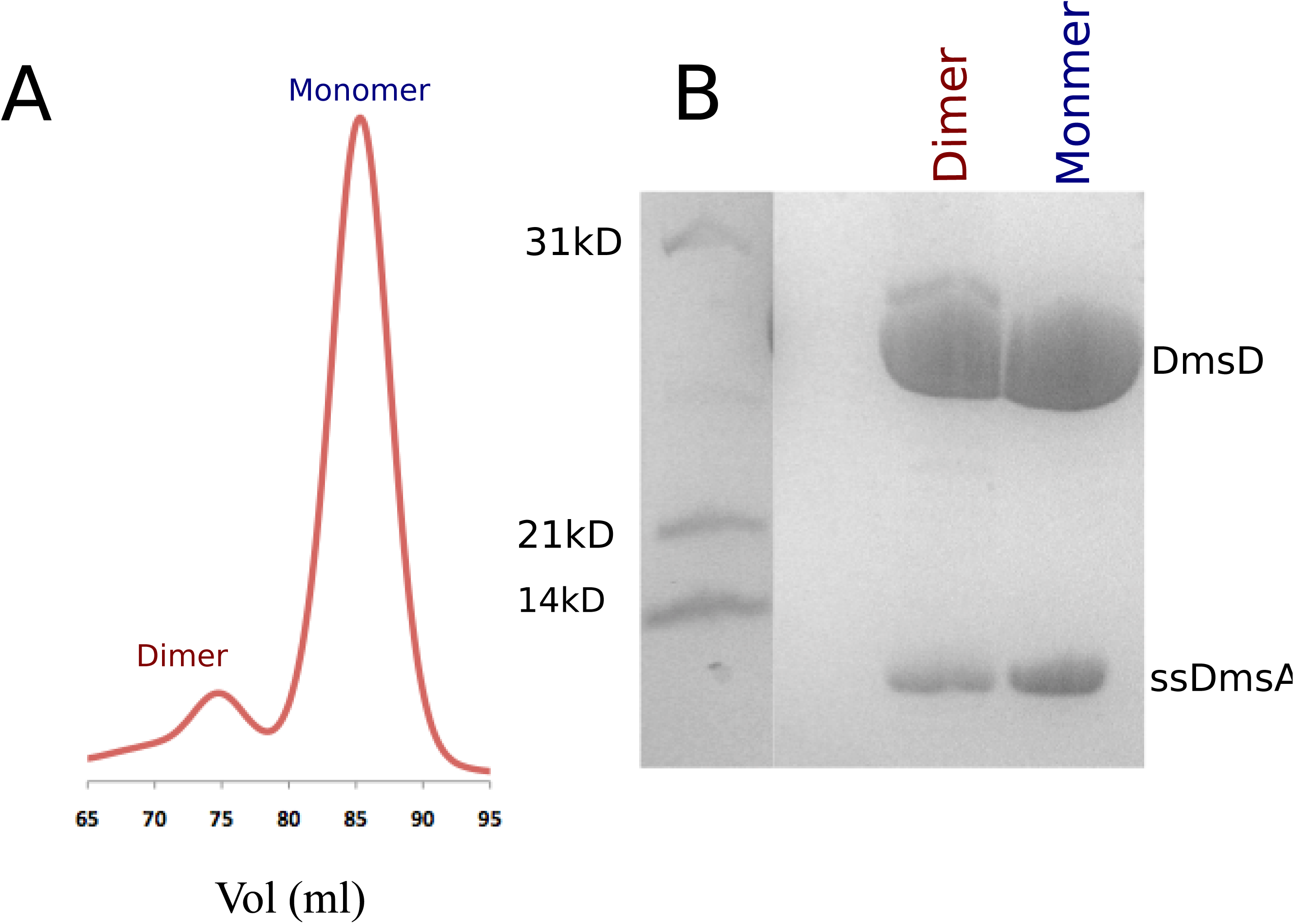

We suspected that a signal sequence bound to DmsD would have a stabilizing effect on DmsD. We tested this using circular dichroism (CD) where we generated melting curves for DmsD alone and bound to ssDmsA (monomeric form) (Figure 4A). The difference in the thermal stability were shown in the (Figure 4B). The thermal stability analysis showed that DmsD is less stable in the presence of ssDmsA.

**Figure 4.**
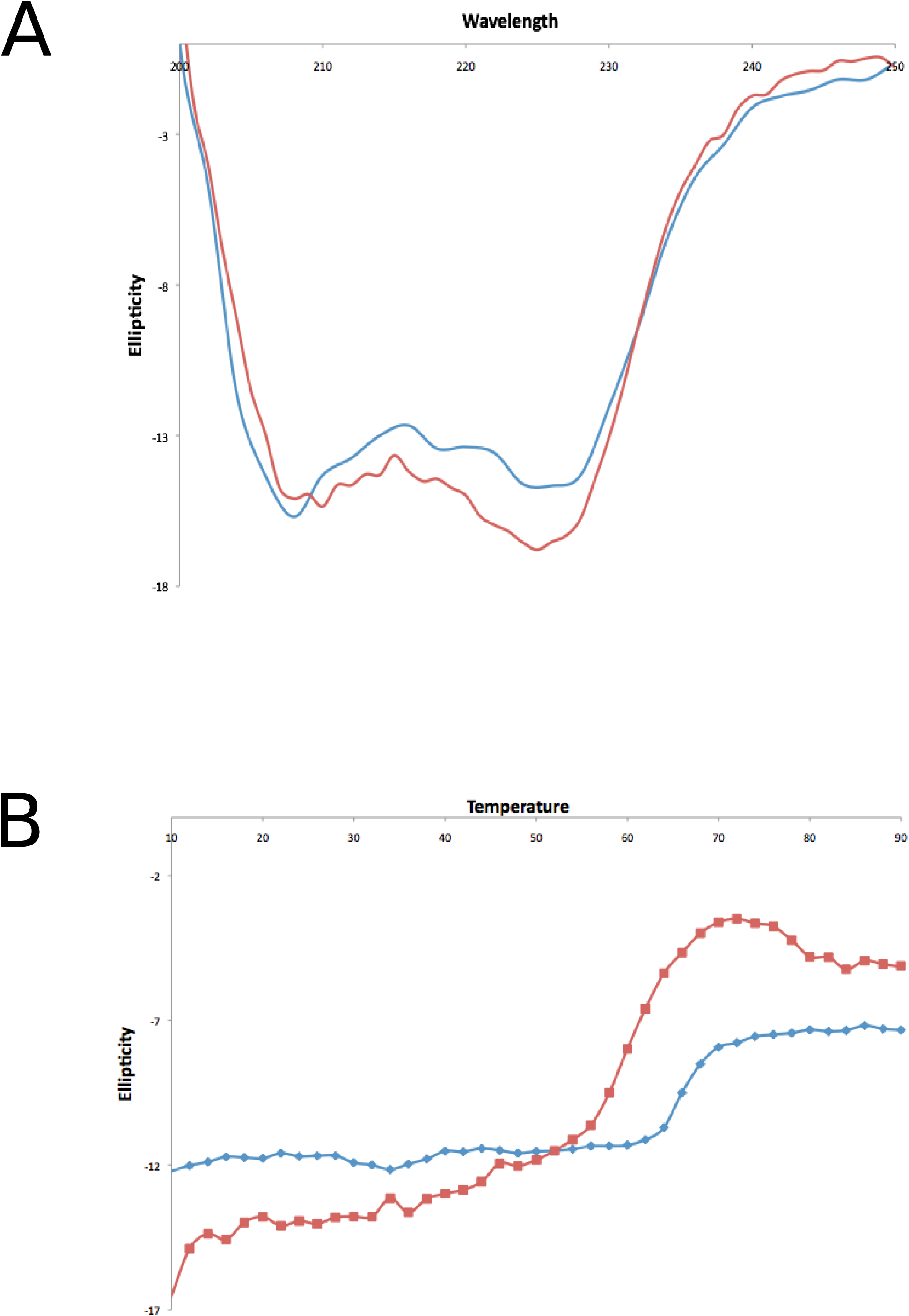

### Tetrameric complex formation

Interaction between TatBC with DmsA, DmsD and DmsD/ssDmsA complex was observed after incubating each component for 2 hrs at 277K and then affinity capture using the 6xHis tag of ssDmsA or DmsD. As a control, TatBC was overexpressed without a tag and ssDmsA/DmsD complex was intact in the presence of detergents (data not shown). Complex formation was confirmed by a shift in the SEC elution profile (Figure 5). The tetrameric complex of DmsA/ssDmsA/TatBC only formed if the monomeric complex DmsD/ssDmsA was used. The dimeric form ssDmsA/DmsD was unable to capture TatBC.

**Figure 5.**
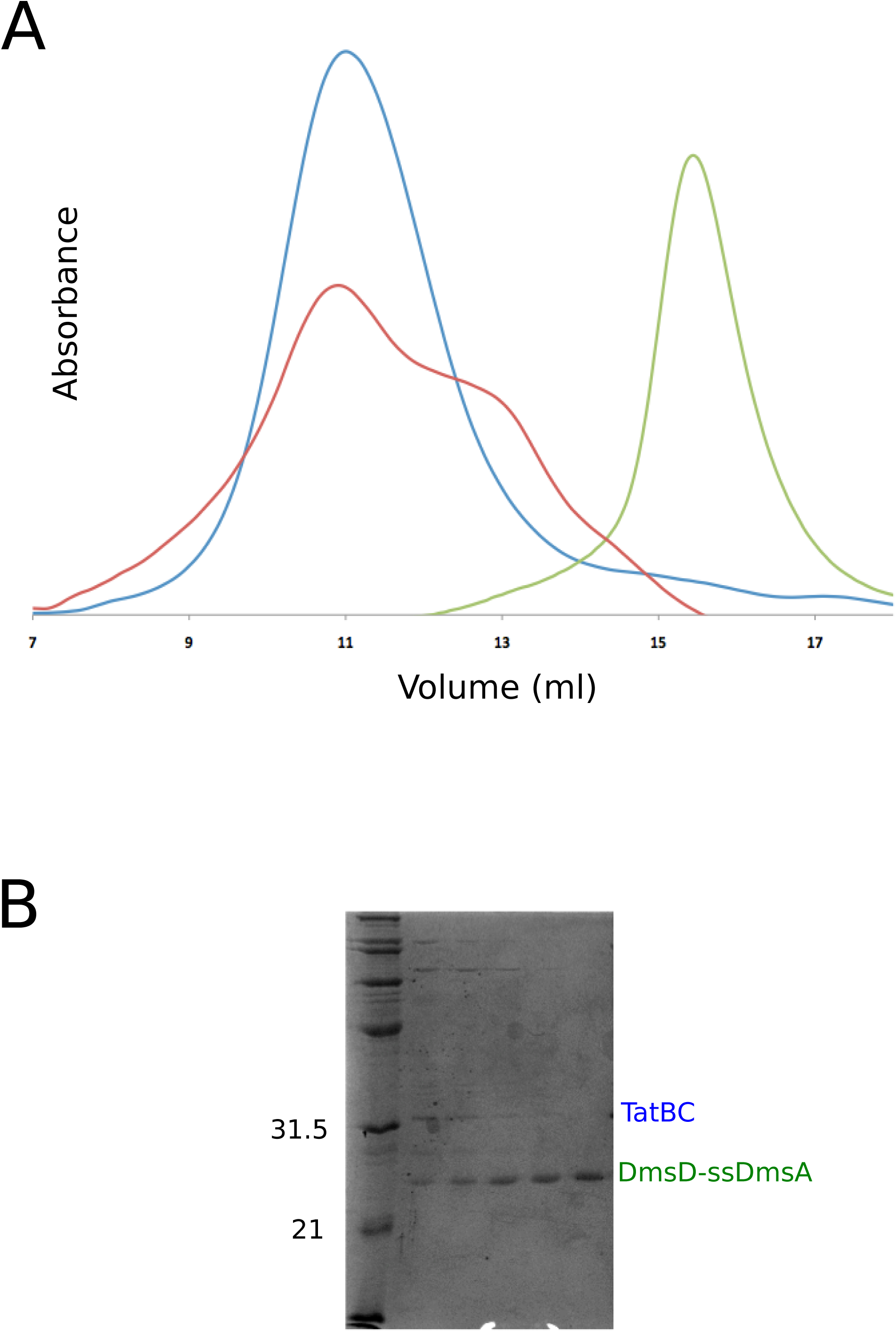

### Sequence comparison of Tat signals with reference to DmsA

Tat signal sequences are typically 30-50 amino acid long (36). The signal sequences of the closely related DmsA and TorA were aligned (Figure 6A). DmsA signal sequences are ∼50 amino acid long while TorA signal sequences are relatively shorter, ∼35 amino acid. An alignment of all Tat signals are recognized by REMP chaperones from *E. coli* is shown in Figure 6B. The alignment shows the conservation of Tat signal motif, located in the border of n- and h-regions, between the various Tat substrate from *E*.*coli*. The h-region is relatively longer in the case of DmsA signal peptide in and Ala-29, Ala-31, Leu-37, Pro-38 and Phe-39 in h-regions are conserved at least among DmsA signal peptides.

**Figure 6.**
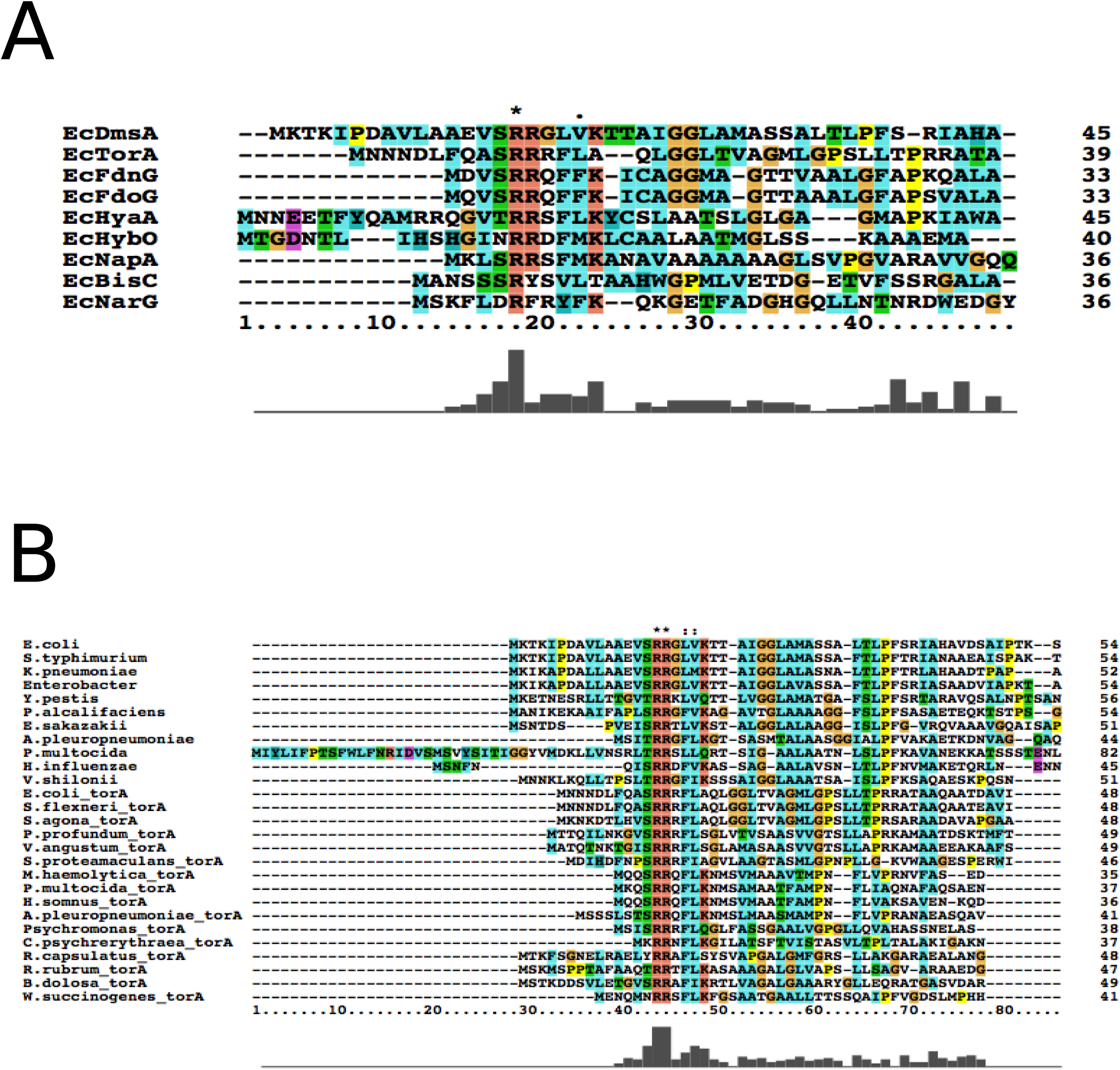

### Effect of truncation of ssDmsA on DmsD and TatC binding

N- and C-terminal truncation mutants of ssDmsA were tested for binding to DmsD. Truncation constructs removed five amino acids at a time from either end of the signal sequence (Figure 1C). In the first round of truncation only C-terminal or N-terminal truncation were tested. Based on these results, double mutants were tested in subsequent cycle. They were tested for binding to DmsD in both *in vivo* and *in vitro* conditions. The results where shown in Figure 7. The deletion of first 10 residues in N-terminal had no significant effect on DmsD binding, except the change in the ratios of the monomeric versus dimeric complex formation compared to wild type ssDmsA (Figure 7B). Ten amino acids could be deleted from the C-terminus before a significant loss of binding occured. Altogether, binding to DmsD required a minimum of 29 amino acids in the signal sequence from residue 13 to 41 (Figure 1C).

**Figure 7.**
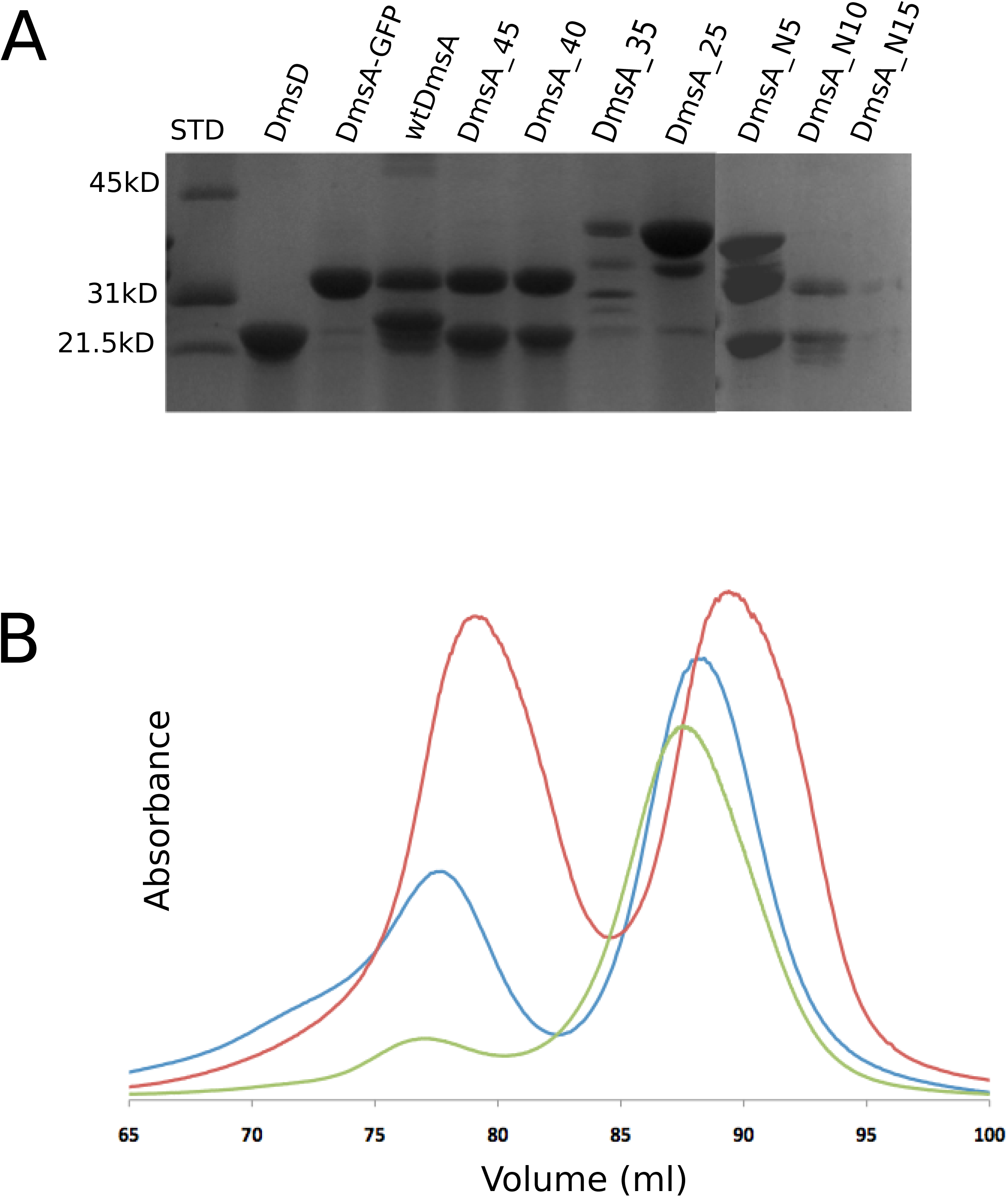

Then all the truncated constructs of ssDmsA-GFP have been tested for its ability to form complex with *Ec*TatBC. Pull down the *Ec*TatBC complex with purified tagged protein of ssDmsA-GFP to determine and map the region in signal peptide responsible for this tetrameric complex formation. From the figure 8 we can clearly observe that there is different forms of complex formation between the *Ec*TatBC and NΔssDmsA/DmsD. The residue deletion from C-terminal of ssDmsA-GFP signal peptide has no effect on the complex formation.

**Figure 8.**
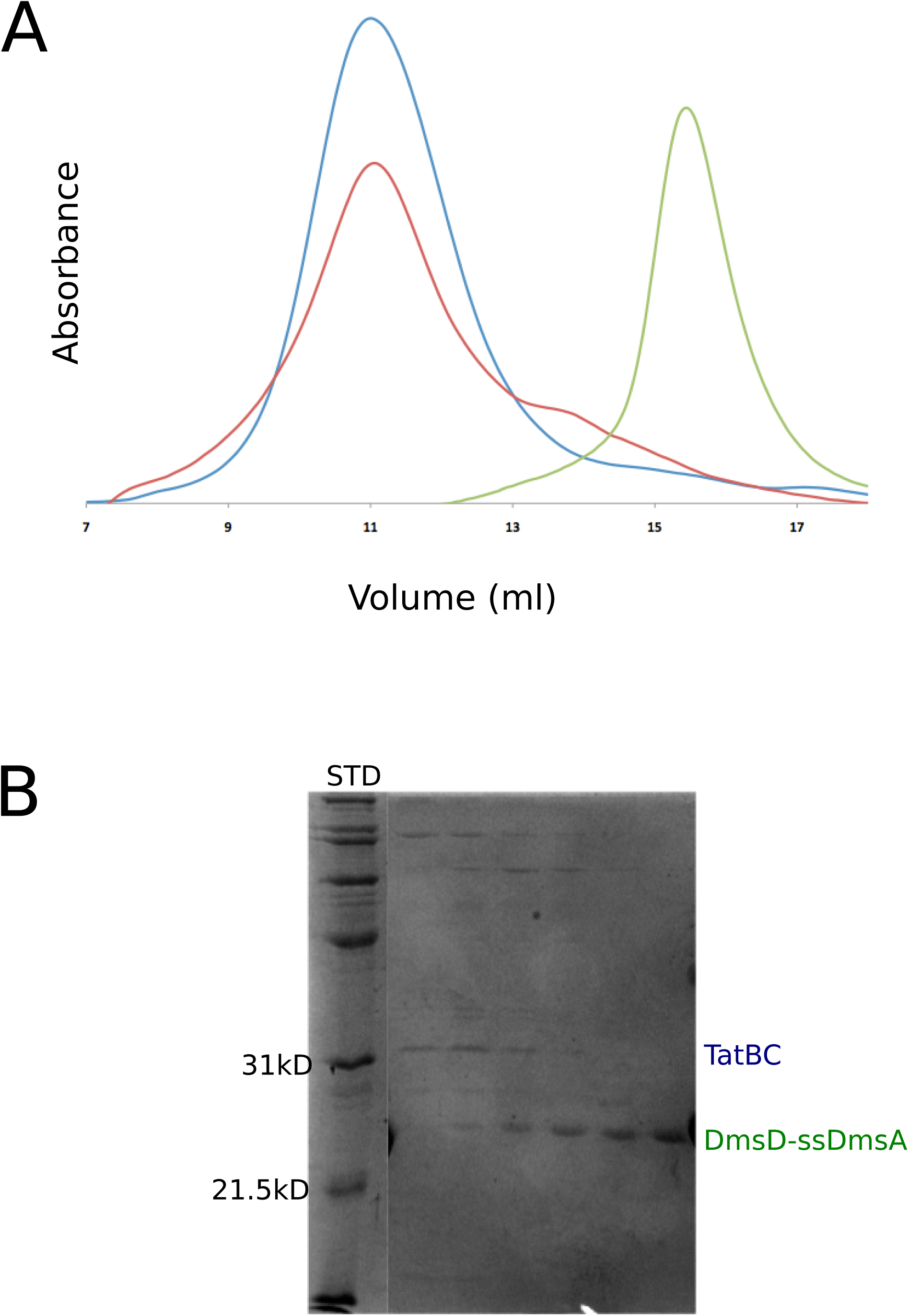

### Sequence determinants of the minimal signal sequence

Alanine scanning of the minimal 29 amino acid long DmsA signal sequence was used to establish the important recognition elements for DmsD binding. Each amino acid was individually replaced with either Ala or Ser (in the case of Ala). We assayed each mutant for the ability to form complex by co-expression and then affinity capture on a Ni-resin. ssDmsA-GFP-6xHis construct co-expressed with untagged DmsD. Using this construct, the visualization of GFP fluorescence allowed for quantification of the expression level of each mutant indicating that they was no significant effect on expression (Figure 9). Additionally, the effect was confirmed *invitro* by mixing saturated amount of purified DmsD with mutants of the ssDmsA-6XHis and followed by affinity purification. The presence of a stable complex is indicated by the ability to capture DmsD visualized by SDS-PAGE (Figure S4). Based on this analysis, Lys 21, Gly27, Ala29, Ala34, Thr36, Leu37 and Phe 39 were all required for complex formation. Mutation of the invariant RR motif completely abolishes the Tat-specific export (39) but had no effect on DmsD binding.

**Figure 9.**
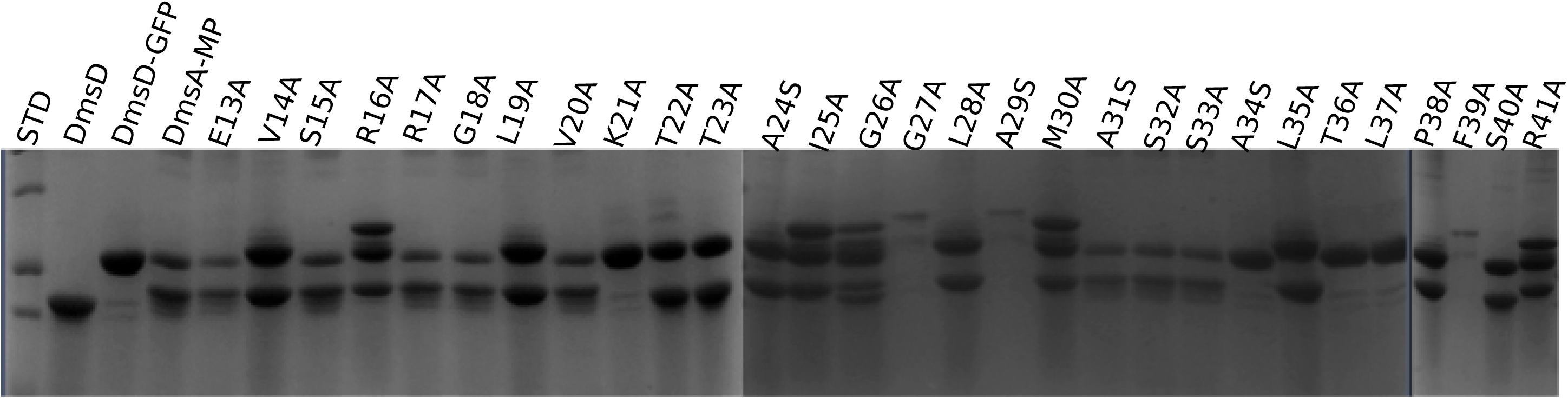

### Kinetic analysis of DmsD/ssDmsA complex formation and interactions with TatBC

We used surface plasmon resonance (SPR) to obtain association and dissociation rate constants during complex formation between DmsD and ssDmsA along with the higher order TatBC interaction. Kinetic rate constants for the binding of the Tat signal peptide of DmsA to DmsD were determined by flowing ssDmsA-GFP to a chip containing immobilized DmsD. As a control, free GFP showed no interaction with immobilized DmsD indicating that any interaction was due solely to the signal sequence (Figure S5).

The key to a successful SPR experiment is the ability to saturate the surface of the chip with the Ligand. We were unable to immobilize the DmsD effectively by affinity tag; therefore, DmsD was attached to the chip by direct amine coupling. We tested a variety of conditions and were able to regenerate the chip using 150mM Nacl and 100 mM glycin buffer pH 9.5. This fully removed the ssDmsA-GFP without disrupting the native conformation of DmsD. These conditions allowed for reproducible binding of ssDmsA to DmsD.

The sensogram of wild type peptide and other truncation mutants were shown (Figure 10). Figure 10 shows the experimental sensogram obtained, association of analyte ssDmsA (wt or mutants) with the DmsD proceeded for 10 minutes and dissociation in analyte free buffer was typically 15 minutes. For each titration, a zero analyte surface as a control was subtracted from test sensogram. The kinetic rate constants (K_a_ and K_d_), as well as equilibrium dissociation constant (K_D_) were estimated by global fitting analysis of the titration curve to the 1:1 (40). The results are summarized in Table 1. Good fitting of experimental data to the calculated curves has been observed, suggesting the correctness of the used fitting model. It provided the insight into the association and dissociation kinetics of the interaction between either wide type or truncated signal peptide of ssDmsA and DmsD. The slower association rate for ssDmsA_wt (K_a_ = 16.4 × 10^4^ Ms-1) and resultant complex between the ssDmsA_wt and DmsD were stable as illustrated by slow dissociation rate (K_d_ =2.2 × 10^−4^s^−1^). Kinetic analysis of the sensogram curve suggests that ssDmsD_45 is more preferred (K_D_ = 55.88 nM) over other truncated forms. But the stable associations were observed upto 15 amino acid deletions, with considerable decrease in affinity. N-terminal truncations had no significant effect on DmsD binding and consistent with the earlier results. The derived minimal peptide has similar K_a_ and K_d_ value suggesting that the essential regions of the signal sequence for the interaction with DmsD.

**Table: 1.**

| **ssDmsA** | **Pull down**<br>**with DmsD** | **K<sub>a</sub> X 10<sup>4</sup>**<br>**(1/Msec)** | **K<sub>d</sub> X 10<sup>-4</sup>**<br>**(1/sec)** | **K<sub>D</sub> (nM)** |
| --- | --- | --- | --- | --- |
| **wt** | **++** | 16.4 ±0.8 | 2.2±0.08 | 56.25±3.8 |
| **45** | **++** | 11.0 ±1.3 | 2.5±0.3 | 55.88±4.9 |
| **40** | **++** | 13.0 ±1.1 | 2.7±0.09 | 64.5±7.6 |
| **35** | **+** | 13.8 ±2.0 | 18.0±0.8 | 59.1±5.7 |
| **30** | **-** | 4.5±0.3 | 51.0±1.4 | 82.0±5.6 |
| **25** | **-** | 5.6±0.7 | 57.7±1.9 | 195.4±12 |
| **Ntrun5** | **++** | 16.3 ± 0.4 | 3.4 ± 0.6 | 58.30± 2.8 |
| **Ntrun10** | **++** | 12.8 ± 0.8 | 4.2 ± 0.2 | 55.98 ± 3.4 |
| **Ntrun15** | **+** | 3.8±0.3 | 23.0±1.1 | 166.9±9.7 |
| **DmsD_Mi**<br>**nimum**<br>**peptide** | **-** | 10.7±0.5 | 2.4±0.1 | 56.51±2.9 |

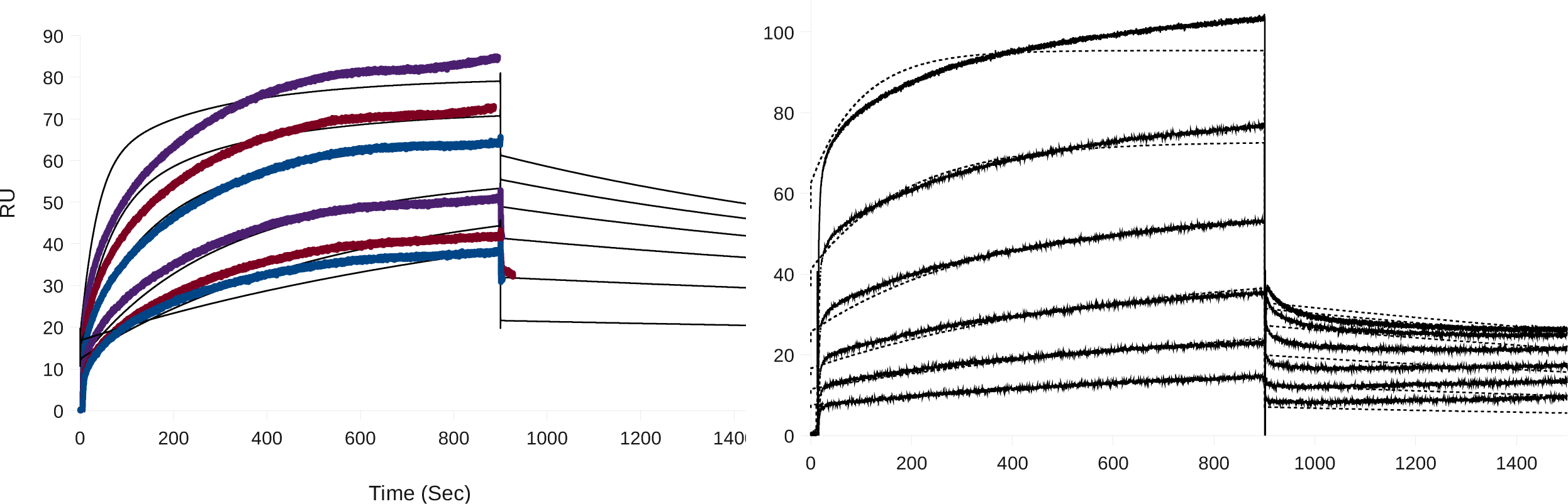

**Figure 11.**
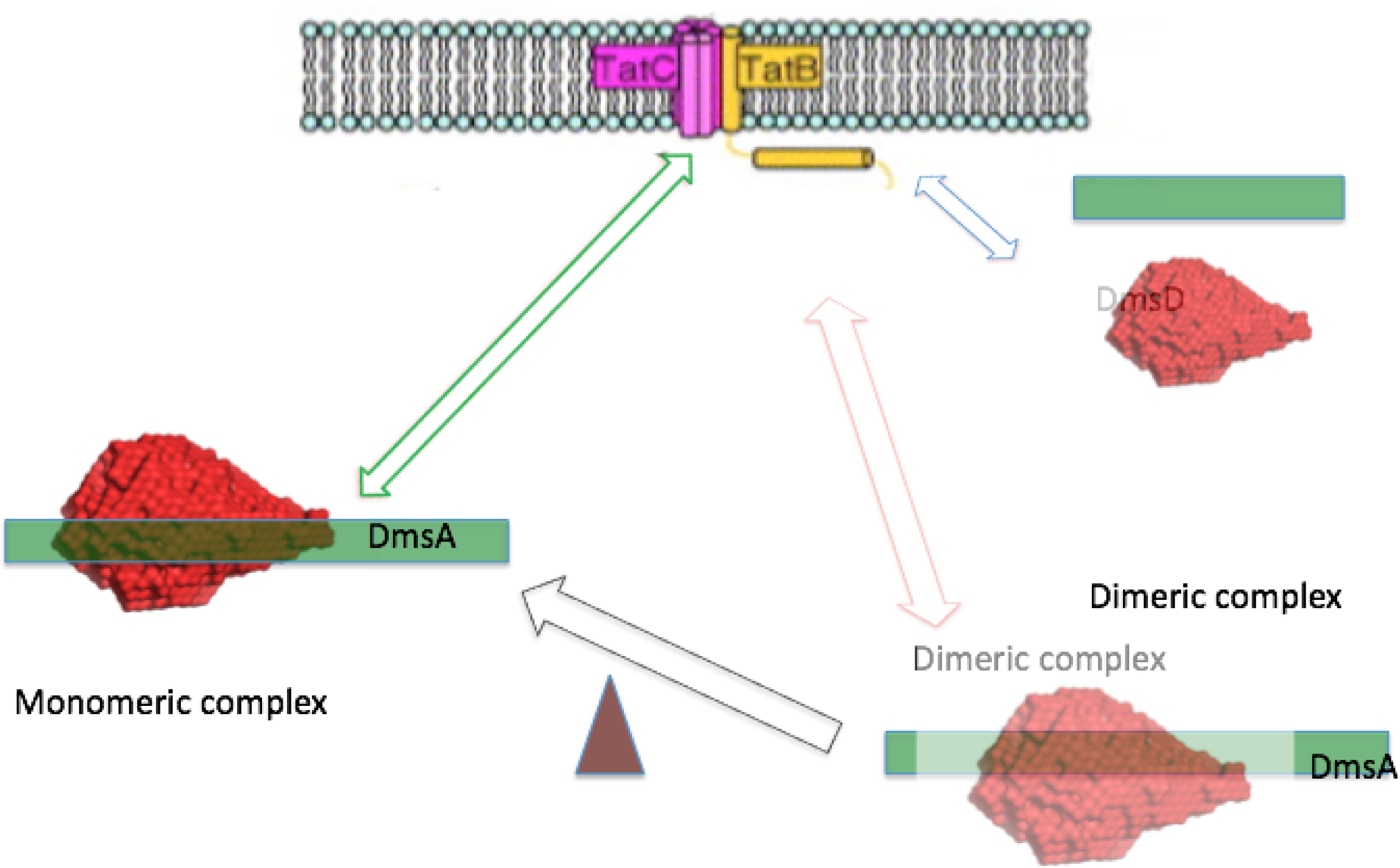

To analyze the interaction of ssDmsA and DmsD with TatBC by SPR, we immobilized TatBC from *E*.*coli*. (Table 2). These interactions were also observed in affinity capture experiments (Figure S6). The affinity constants were higher for the DmsD/ssDmsA complex compared to the individual components. Interestingly, although the N-terminal truncation had no effect on the DmsD/ssDmsA complex formation, truncations significantly lowered the affinity towards TatBC suggesting a role in the higher order complex formation. Moreover, this effect was even seen in measurements using ssDmsA alone. It is worth to note that the derived minimal peptide has significantly less affinity towards the TatBC complex. Even though minimal peptide has showed no difference compared to wild type ssDmsA in the formation of ssDmsA/DmsD complex. This result denotes that N-terminal of signal peptide may be involved in the interaction with translocan complex. Another striking observation is that dimeric form of DmsD/ssDmsA complex has less affinity towards TatBC compared to monomeric form which is on par with the result of affinity capture experiments.

**Table 2.**

| **ssDmsA or and**<br>**DmsD** | **K<sub>a</sub> X 10<sup>4</sup>**<br>**(1/Msec)** | **K<sub>d</sub> X 10<sup>-4</sup>**<br>**(1/sec)** | **K<sub>D</sub> (μM)** |
| --- | --- | --- | --- |
| **ssDmsA_wt** | 9.0 | 5.2 | 5.8 |
| **ssDmsA_45** | 12.3 | 3.4 | 5.3 |
| **ssDmsA_40** | 15.2 | 4.2 | 6.8 |
| **ssDmsA_35** | 20.8 | 1.6 | 8.2 |
| **ssDmsA_30** | ND | ND | 11.4 |
| **ssDmsA_25** | ND | ND | 10.4 |
| **ssDmsA_Ntrun5** | ND | ND | 7.3 |
| **ssDmsA_Ntrun10** | 30.7 | 2.3 | 15.6 |
| **ssDmsA_Ntrun15** | ND | ND | 18.7 |
| **DmsA_Minimum**<br>**peptide** | 26.5 | 1.7 | 19.8 |
| **ssDmsAwt RR**<br>**mutants** | ND | ND | 8.8 |
| **DmsD alone** | 1.1 | 3.1 | 2.8 |
| **ssDmsD/DmsD**<br>**monomer** | 3.0 | 5.1 | 1.7 |
| **ssDmsD/DmsD**<br>**Dimer** | ND | ND | 13.2 |

## DISCUSSION

This work reveals that DmsD binds to the N-terminal signal peptide of ssDmsA and forms two distinct populations. The results from the Biacore analysis and the SEC data or affinity capture were well correlated. Based on this data we have designed the minimum peptide for formation of complex, which consist of 29 amino acids. And the kinetic analysis from SPR suggest that this peptide is having faster association rate (K_a_ = 10.7 × 10^4^ Ms^−1^) compare to all other mutant and wild type, and the affinity is very close to the wild type peptide K_D_ = 56.5 nM.

The sequence comparisons of all Tat signal, which are having binding chaperons reveals some region are more conserved (Figure 6B). In spite of the divergence in Sec and Tat transport systems, they do have features in common. The signal peptides of both pathways are evolutionarily well conserved and display the same tripartite organization. Even though overall conservation signal peptide found in Tat substrate do have several exclusive features as compared to those found in Sec substrates. Tat substrate signal n-region is much longer, and they have twin arginine motif in the n- and h-region boundary. The h-region in Tat substrate usually longer and also has less number of hydrophobic residues compare to Sec signal. An additional lysine or arginine is often present in the c-region of Tat signal peptide, which serve as sec avoidance motif (41).

The sequence alignment provides some inferences with regard to the mechanism of substrate specificity by REMPs. Homology of the entire REMP sequences and the hydrophobic regions of signal peptide imply recognition specificity, with importance on the position and length of the continuous hydrophobic stretch following the Tat motif. The varying architecture of this hydrophobic stretch is likely to adapt to the binding pocket of different REMP structural classess such as those recently described (25). The higher percentage of sequence conservation of DmsD/HybE than for observed NapD/HybE pairs suggest that homology of REMPs is not the only factor consider whether two REMPs will interact with the same signal peptide (42).

The truncation mutations of ssDmsA reveals that the C-terminal part of the ssDmsA is more directly involved in DmsD binding. In case TorA/ TorD system truncated signal peptide of 10-36 of TorA essentially identical to full length and showed tighter binding (35). Also showed that there is no preference over twin-lysine peptide, which suggest that twin arginine motif is not essential for chaperone recognition (35). This is on par with our Ala scanning results where the replacement of RR with Ala has no effect on the DmsD binding (Figure 9). The another study suggested that Ser in −1 position and Leu in +2 position in the peptide were essential for the translocation (43).

A leucine rich region within the signal peptide h-region is shown to be involved in TorD binding both *in vivo* and *in vitro* (36), which correlates well with our alanine scanning results in ssDmsA/DmsD system (Figure 9). Before the pre-protein interacts with Tat translocon it interact with inner membrane that is stabilized by both electrostatic and hydrophobic contribution (44). The studies are revealed that the positively charged signal peptide region and anionic lipid head groups play major role in the association of preprotein with the lipid bilayer. And h-region of the signal peptide interacts with the apolar environment of phospolipid (44). It has been suggested that increasing hydrophobicity of h-region favors the membrane binding rather chaperone binding.

Here we report for the first time that the DmsD/ssDmsA interaction analysis showed undoubtedly the presence of two forms of complex. And equilibrium of these forms can be modulated by N-terminal truncation. Further *in vitro* analysis showed that the concentration of DmsD also influence the architecture of the complex

The NarJ/NarG system, which is similar to DmsA/DmsD has been investigated thoroughly in structural level by NMR. This result suggests that NarG signal peptide is in helical conformation and also there is change in conformation in NarJ upon the signal peptide binding. pH depended modulation of the peptide binding affinity has also been observed (45).

Translocation involves a very intimate association of signal peptide with the receptor complex binding site (46). Previous reports suggested that DmsD alone interact with Tat apparatus especially TatB and TatC (47). Here we could able isolate and identify the regions responsible for this interaction between TatBC with DmsD, ssDmsA and ssDmsA/DmsD complex. The N-terminal of the signal peptide has been mainly involved in the TatBC recognition, which includes the Tat motif. The effect of mutation on the Tat motif has been well studied on the translocation previously (2). The N-terminal of Tat motif has seems to play some role in the complex formation. Even though the deletion of the N-terminal upstream of the Tat motif still forms complex with TatBC, the association is weak and forms different type of complex. The C-terminal of the signal peptide doesn’t seem affect binding.

TatB is found in contact with the entire signal sequence and adjacent parts of mature part of the protein, but in TatC the interaction restricted to a discrete area around the consensus motif (2).

Among monomeric and dimeric types of ssDmsA/DmsD complex, only the monomeric population interacts with TatBC and forms the tetrameric complex. In case of monomeric complex of DmsD/ssDmsA, both interaction site of signal peptide and the DmsD may be still accessible for the Tat translocase assembly. The dimeric complex could be formed due to stable and very strong interaction of DmsD molecules. The DmsD concentration depended binding towards signal peptide has been observed previously (34) and in this work (Figure S1).

It is possible that both of these types of complex may be needed and serve as mechanism for the stringent incorporation of co-factor and quality control of DmsA maturation. The model that emerged from this study has been represented in Figure 12. After the nascent peptide of DmsA emerge from ribosome DmsD form dimeric complex. So the DmsA signal has been complexly masked and it is inaccessible for the translocon and also this dimeric complex obstructs the DmsD interaction with the receptor complex. Once the cofactor has been inserted and after completion of sub-unit assembly the DmsD dissociate to monomeric form which has now exposed all its interaction sites over signal peptide as well as in DmsD and this monomeric complex readily interact with translocase assembly. There is enough evidence that the mature part of DmsA interact with DmsD (48). The other possible scenario is DmsD may form monomeric or dimeric complex in normal circumstance but once the degradation increases due to improper folding and assembly (49) it may trigger the DmsD synthesis that eventually increases the relative concentration of DmsD with respect to DmsA and equilibrium shifted towards the dimeric form. Dimeric form restricts the access to the translocon prematurely and leads to stringent quality control of DmsA maturity as a functional assembly. One interesting and supporting fact to this model has been observed in this studies was once the N-terminal of the signal peptide truncated there is shift in monomeric and dimeric form of the complex which also indirectly tells us the steric hindrance may be playing role in the complex regulation.

