## Supplementary figures and images for "Defining the DmsA Signal Sequence Interaction with DmsD And TatBC"

### Supplement Figures

Figure S1

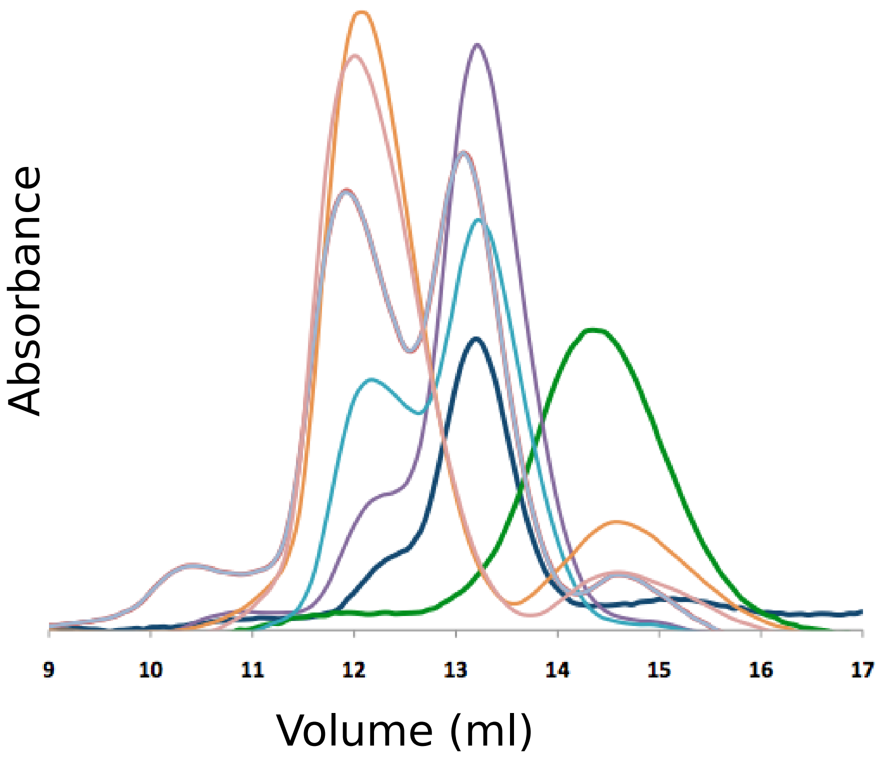

Figure S2

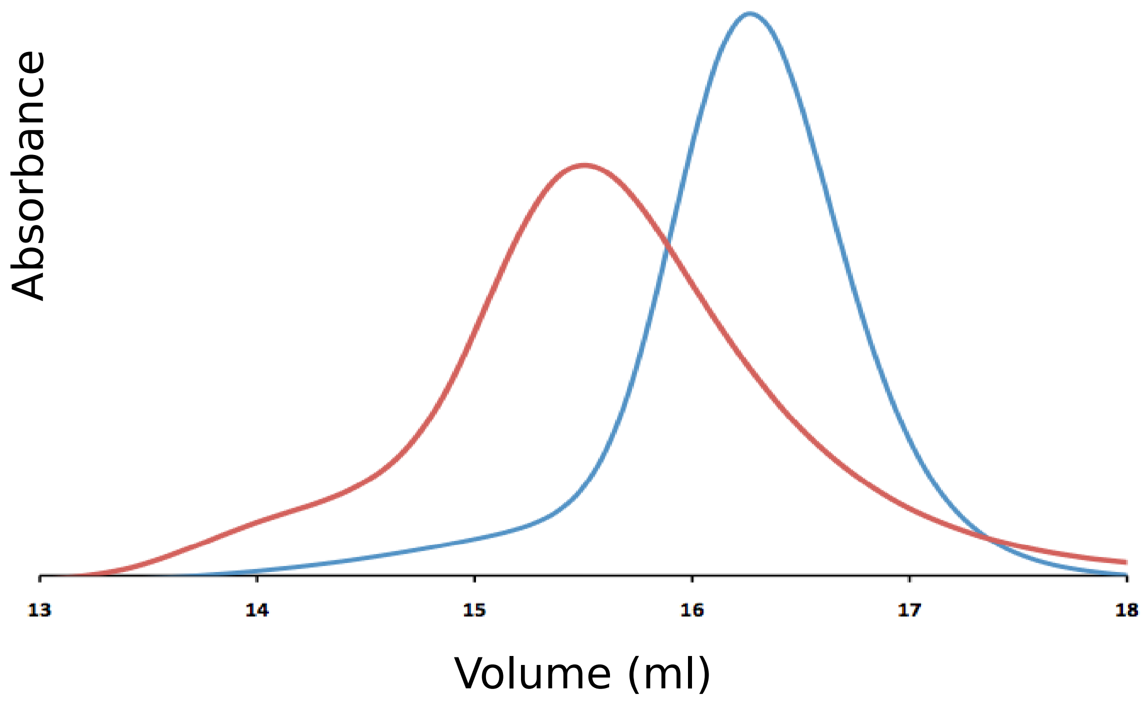

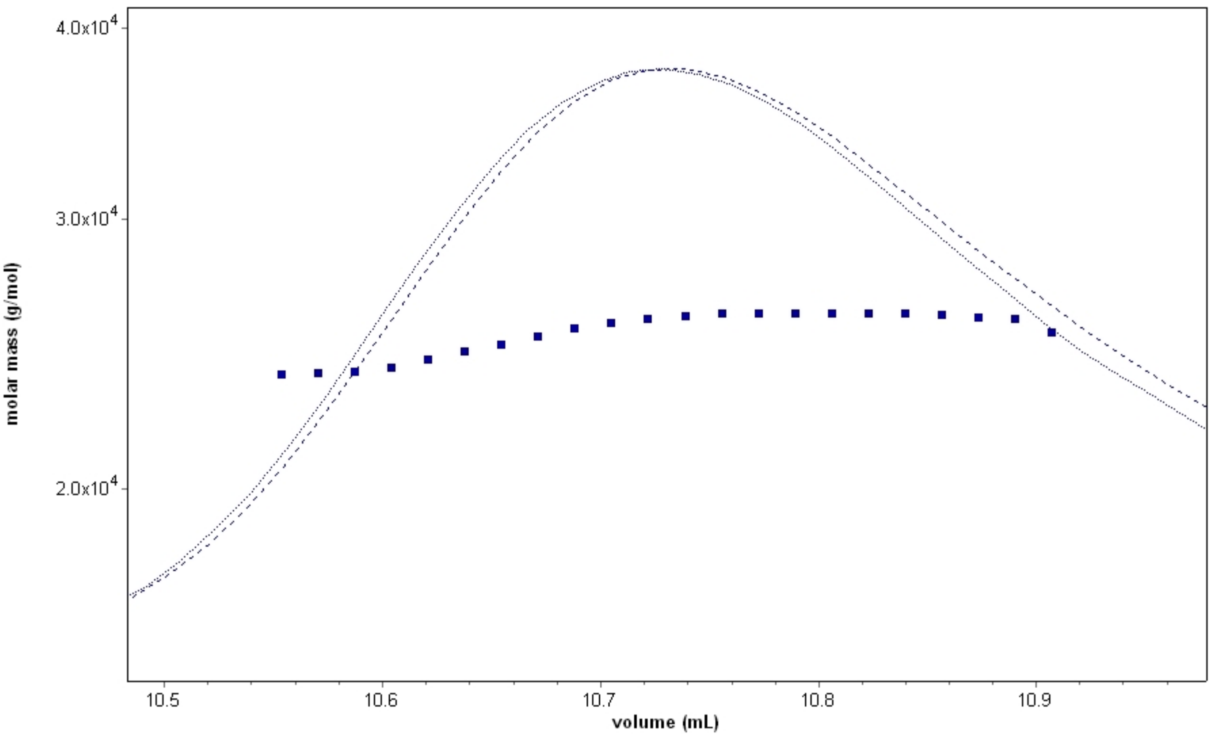

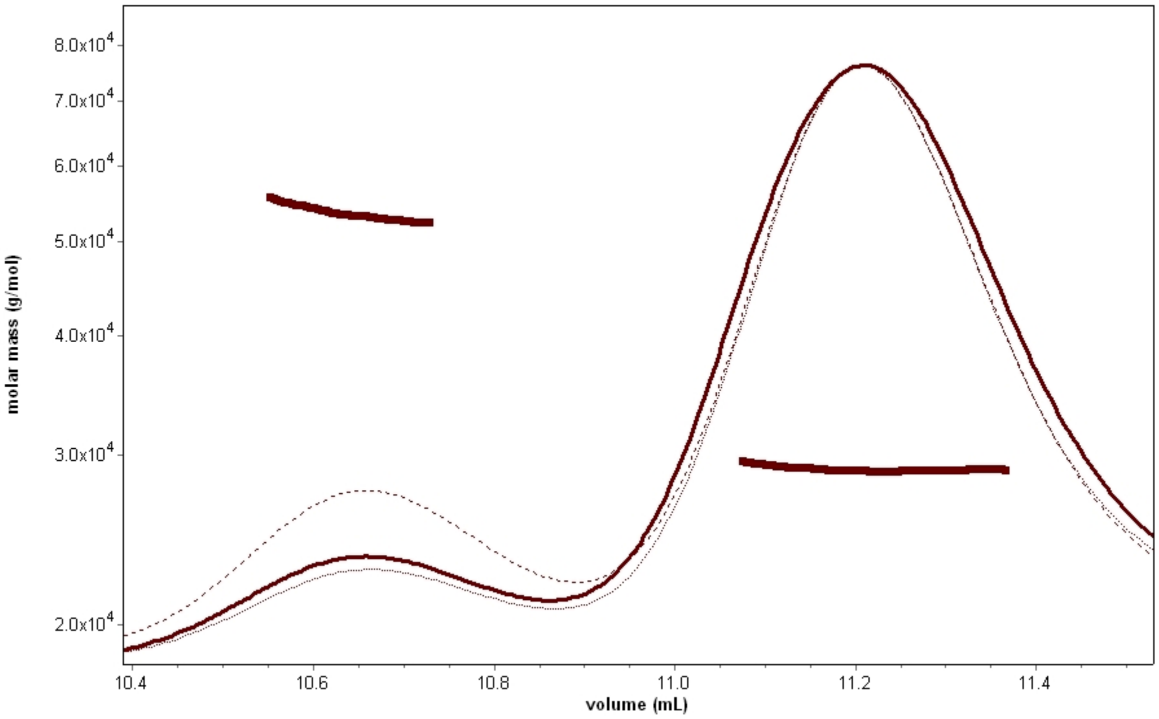

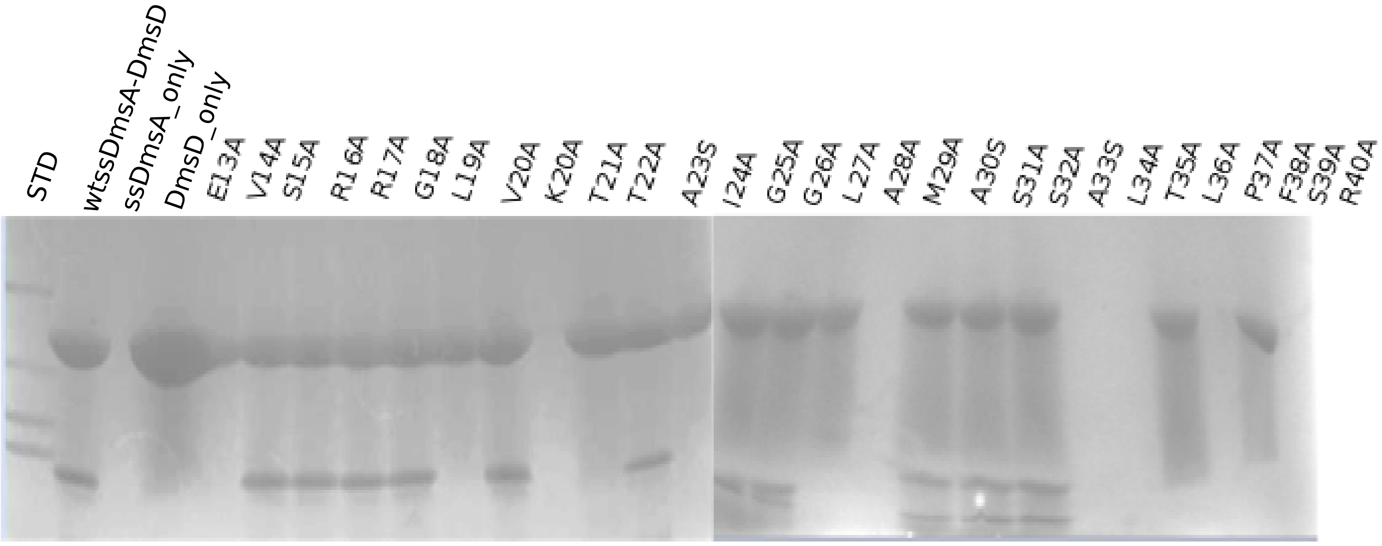

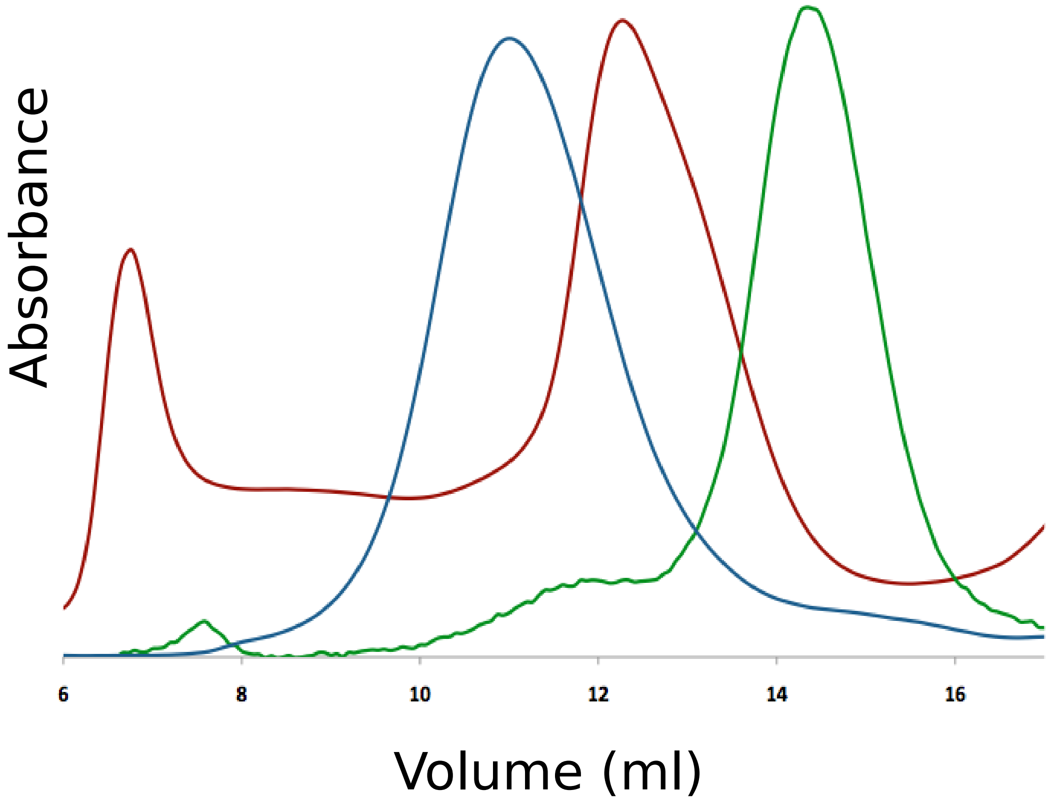

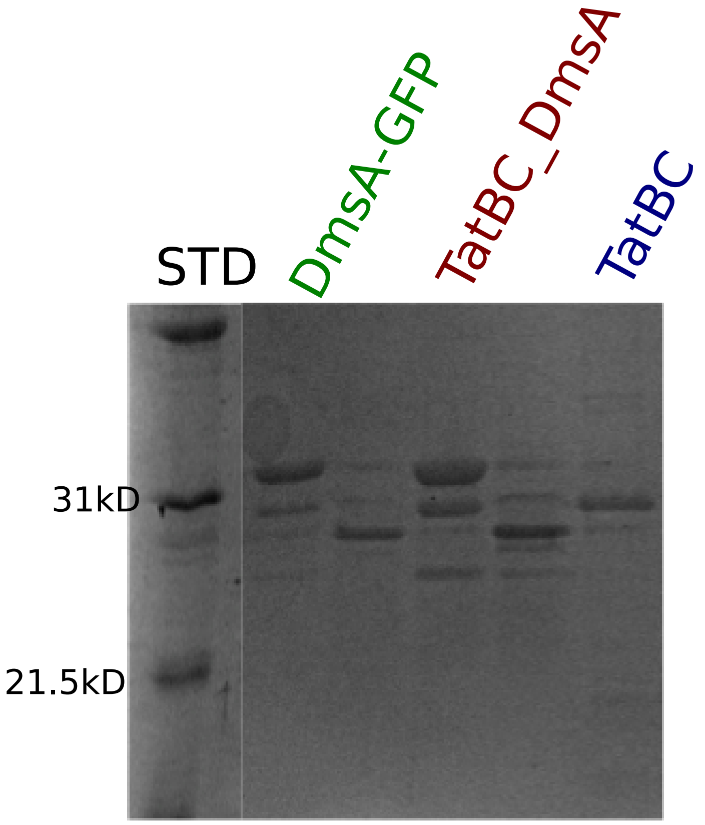
